# Scanning Electron Microscopy Study of Bacterial Growth in Mycelial Extracellular Matrices

**DOI:** 10.1101/2025.02.12.637402

**Authors:** Davin Browner, Andrew Adamatzky

## Abstract

Fungi and bacteria are found living in a wide variety of environments and their interactions are important in many processes including soil health, human and animal physiology and in biotechnological applications. The specificity of interaction between these microorganisms in co-culture is difficult to establish. For example, differentiation between trivial processes as a result of stochastic mixing compared with mutualistic or antagonistic interactions. Here, we investigate a single morphological feature of co-cultures of planktonic bacterial growth within biofilm-forming liquid cultures of mycelium. Namely, the attachment of bacterial co-habitants of species *Bacillus subtilis* to fungal hyphae of species *Hericium erinaceus*. The bacteria-in-mycelial-biofilm method was developed and utilised to allow for attachment of bacteria to hyphae via containment within extracellular polymeric substances (EPS) and the overall extracellular matrix (ECM) of the mycelium. Attachment structures appear to result from the hyphal surface as a result of production of EPS. The mean biofilm area across T1-3 was 3.90 (*µ*m^2^) ± 0.72 (*µ*m^2^) and the mean percentage coverage was 18.33 (%) ± 5.52 (%). The bacterial biofilm components could not be ruled out as co-contributing to formation of attachment structures due to the structures being present connecting individual bacterium as well as to hyphae.

## 1. Introduction

Bacterial microorganisms occur in almost all ecological settings and their ubiquitous presence has led to antagonistic and mutualistic relationships with fungi. These interactions are important components in a large number of environmental processes [1, 2]. Co-communities have been described to exist in virtually all ecosystems and across a wide diversity of fungal and bacterial families. In artificial settings, fungal and bacterial communities of microorganisms have been used in food production and agriculture [3], pharmacological [4], bioremediation [5] and biotechnological applications [6]. When directly observed, the bacterial and fungal interactions appear to intrinsically modulate the behaviour of either or both of the microorganisms, for instance in reduction of fungal growth rates due to antagonistic bacterial co-habitants in contaminated fungal cultures. However, prediction or even identification of this modulation capacity and interaction is difficult based on existing knowledge of the biology of the isolated microorganisms grown in pure cultures. The integration of fungal and bacterial components is complicated by different levels and specificity of interactions [7]. For example, stochastic mixing of bacterial components in liquid cultures of fungi is unlikely to be representative of any actual causal relationship [7]. In contrast, co-occurrence can lead to integrated biophysical and metabolic interactions.

As a result, interactions can be simple, specific or trivial [7]. Highly specific interactions have been observed for endofungal bacteria [8] and in early emerging bacteria [9]. The question of whether such cocultures are also co-evolving is complicated by factors such as utilisation of infection or artificial insertion in endofungal studies of bacteria. However, studies with sufficient duration of growth have shown the respective microorganisms independently developing and co-evolving in co-culture. Both intracellular [10] and extracellular [11] relationships have been observed. Supposed interactions exist within a spectrum of activity relating to a variety of processes including growth, reproduction, transport,movement,nutrition, stress resistance and pathogenicity [7]. These processes impact the microorganisms at different levels and with ranging specificity resulting from the combined physical associations (biofilm, free cells, intracellular), the molecular dialogue between the organisms (direct or indirect) and the environmental conditions and/or host activity [7].

Biofilms may play an important role in mediating fungal-bacterial co-culturing including facilitating antagonistic, mutualistic or inconsequential interactions. Bacterial and mycelial biofilms are both independently known to adhere strongly to various surfaces [12] [13]. Potential interactions in co-cultures may proceed following the attachment of bacteria to hyphae via containment within the extracellular polymeric substances (EPS) and macroscale extracellular matrix (ECM) of the fungal species. Conversely, bacteria may selectively proliferate in certain advantageous regions of the fungal culture and attach to fungal hosts. Many biotechnological processes rely on the formation of bacteria co-habitants in liquid cultured mycelia including fermentation and brewing [14], cheese ripening [15], bioremediation [16] and natural product discovery and synthetic biology [17]. In non-sterile artificial conditions contamination of fungal liquid cultures is a common occurrence in production of mycelium and fungal fruiting bodies. Bacteria have received attention in fruiting body cultivation in a wide number of species of fungi for potential mutualistic relationships [18].

The properties of bacteria growth within mycelial biofilms, how commonly such growth might occur and the various resulting interspecies and inter-kingdom interactions are underexplored. Apart from in endofungal co-cultures, the successful introduction of extracellular bacteria appears to primarily depend on the level of biofilm rejection or integration. Scanning electron microscopy (SEM) studies of the morphology of co-cultures of bacteria and fungi can be used to investigate the specificity of interactions. In particular, SEM can be used to identify whether attachment or other complex morphological growth patterns have occurred. In this paper, we outline investigation of the morphology of bacteria grown inside mycelial biofilms in the form of overnight cultures of bacteria *Bacillus subtilis* and fungal co-habitant *Hericium erinaceus. B. subtilis*, known also as the hay bacillus or grass bacillus, is a gram-positive, catalase-positive bacterium, found in soil and the gastrointestinal tract of ruminants, humans and marine sponges. The imaged structures repeatedly show bacterial attachment to the hyphal surface via polymeric substances. We plan to use these structures to investigate the modulation capacity of bacteria on mycelia in terms of extracellular electrical activity. Therefore, the focus of this study is on establishing that bacteria can attach to mycelial hyphae. Attachment structures appear to result from the hyphal surface as a result of production of EPS.

## 2. Methods

### 2.1. Co-culturing

Mycelium of *H. erinaceus* is readily cultivated in liquid culture, agar or agar slants. The basidiomycetes species *H. erinaceus* can be grown via liquid static fermentation. This method produces a large amount of extracellular materials including polysaccharides and proteins that contribute to the formation of ECMs. Mycelial cultures of *H. erinaceus* were sampled from stocks previously identified as showing biofilm-forming capacity in liquid media and co-culturing with bacterial species. Using these stocks liquid cultures were grown from agar plate samples in malt liquid culture media (1.5 %). Fungal biofilms were grown in submerged cultures within glass vials sealed with butyl rubber stoppers and incubated at 25 C^°^ for 24 hours. Then the liquid cultures were grown at room temperature (20^°^C) for 4 days in 20 ml vials without agitation. Lack of supplementation simplified the resulting SEM analysis due to removal of impurities and trace contents during dehydration. The vials were enclosed by self-healing butyl-rubber stoppers which functioned as a sterile method to initiate growth via injection of liquid cultures and restricted oxygen supply to increase fermentation and EPS formation. Mycelial growth occurred in an enclosed growth tent and without exposure to light. Temperature fluctuations were avoided by use of a temperature controlled lab set to (20^°^C). Bacterial species were grown overnight in an incubator at 35^°^C from stocks of *B. subtilis* (LZB 025). The resulting bacterial cultures were then transferred to the vial containing the mycelial liquid cultures and the vial was tilted and left at room temperature in a dark environment for 12 hours. Solutions of 100 *µ*L of bacterial media were transferred into each of the triplicate samples. Following this resting period the co-culture samples underwent further processing for SEM imaging as detailed below.

### 2.2. Sample preparation for imaging co-cultures

Using sterile techniques and a dissecting microscope, the co-cultures were transferred from the growth vial to an empty 20 ml glass vial. The samples were fixed in 4% glutaraldehyde in PBS for 1 hour at room temperature (20^°^C) and then rinsed 3 times with PBS and stored until required.

### 2.3. Dehydration using hexamethyldisilazane

Fixed co-cultures were dehydrated in a graded ethanol series. Ethanol was replaced by hexamethyldisilazane (HMDS) in 5-minute steps in the following ratios: 1:2, 1:1 and 2:1. This was followed by two rinses in 100% HMDS. The co-cultures were left to dry overnight on filter paper in a sealed Petri dish and mounted on SEM stubs. They were sputter-coated with gold immediately prior to imaging. They were then imaged using the scanning electron microscope. More than two dehydration steps were found to dehydrate the samples resulting in complete collapse of the hyphae.

### 2.4. Scanning electron microscopy (SEM) preparation, parameters and image processing

The FEI Quanta 650 FEG scanning electron microscope (SEM) (FEI Company, U.S.) was used to image the samples and produce micrographs. Image processing and analysis was performed using ImageJ [19]. The scale was set using the metadata from the scanning electron microscope and each sample. Measurements were then performed taking into account the 3D structure of samples. Occluded, deformed or partially visible bacteria and hyphae were not included in measurements.

## 3. Results

The mean length of co-cultured bacteria in the triplicate samples was 1.4 *µ*m ± 0.4 *µ*m. The smallest sized bacteria was 0.53 *µ*m and the largest observed was 2.6 *µ*m. The mean width of bacteria was 0.5 *µ*m ± 0.1 *µ*m. The minimum width bacteria was 0.25 *µ*m and the largest was 0.85 *µ*m. Bacteria with longer lengths were likely to be the result of binary fission processes that were ubiquitous in the co-culture samples and indicate active physiological processes occurring prior to fixation. The mean width of hyphae in the cocultures across triplicates was 3 *µ*m ± 0.5 *µ*m compared with a mean width of hyphae in the monocultures of 3.4 *µ*m 0.5 ± 0.4 *µ*m. The largest hyphal width observed for the monocultures was 4.5 *µ*m and the smallest width was 2.5 *µ*m. For the co-cultures the max width hyphae observed was 4.2 *µ*m and smallest 2.3 *µ*m. Using the same methods the attachment structures were measured in higher magnification images (1 *µ*m scalebar). The mean length of attachment structures to hyphae was 0.3 *µ*m ±0.1 *µ*m. Dehydration artifacts were unavoidable and impacted the morphology of hyphae considerably leading to hyphae width estimations on the lower end of typical widths of 1 *µ*m - 10 *µ*m. The procedure seems to have had limited effect on the bacterial co-habitants with only sporadic instances such as burst bacteria apparent in the sample images. The morphological features are detailed for each of the triplicates in Table 1.

**Table 1:** Comparison of basic morphological features of mycelial controls and co-cultures (magnitude of 5 *µ*m scalebar).

| Sample | Mean & Std Number of Bacteria (per $\mu\text{m}^2$) | Mean & Std Length ($\mu\text{m}$) of Bacteria | Mean & Std Width of Bacteria | Mycelial Hyphae Width Mean & Std |
| --- | --- | --- | --- | --- |
| Mycelial controls T1-3 (5 $\mu\text{m}$ scalebar) | $0 \pm 0$ | N/a | N/a | $3.4 \pm 0.5$ |
| Co-cultures T1-3 (5 $\mu\text{m}$ scalebar) | $55.64 \pm 14.22$ | $1.4 \pm 0.4$ | $0.5 \pm 0.1$ | $3 \pm 0.25$ |

The Sato ridge detection algorithm was used to detect ridge-like structures in the SEM images. It is based on analysis of the eigenvalues of the Hessian matrix of the image. The algorithm enhances ridge-like structures while suppressing noise and other structures. The remaining biofilm components appeared as ridge-like structures contrasting hyphae and bacteria.

Given an image *I*, the Hessian matrix *H* at each pixel was defined as:

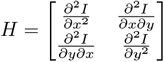

The eigenvalues λ_1_ and λ_2_ of the Hessian matrix were computed. For ridge detection, the following conditions were used:

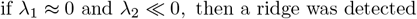

The Sato filter response *R* was given by:

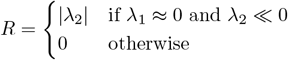

The biofilm area was calculated using custom Python code primarily utilising the scikit-image library (also known as skimage) [20]. The code performed image analysis on the dataset of 1 *µ*m scalebar micrographs to calculate the area and coverage of biofilm in the images. The images were processed to convert them to grayscale, thresholded to identify biofilm components, and then analyzed to compute the area and coverage percentage.

Let *I* be an image with dimensions *M* × *N*. The scale for each image was defined as:

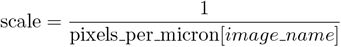

where pixels per micron was a dictionary containing the number of pixels per micron for each image.

The area of vessels *A*_*υ*_ in microns squared was calculated as:

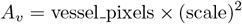

where vessel pixels was the number of pixels identified as vessels.

The total area *A*_*t*_ in microns squared was:

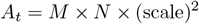

The percentage of vessel coverage *P*_*υ*_ was:

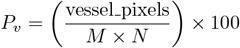

The biofilm areas and percentage coverage are detailed for each of the triplicates in Table 2. The mean biofilm area for T1-3 was 3.90 (*µ*m^2^) ±0.72(*µ*m^2^) and the mean percentage coverage was 18.33 (%) ±5.52(%).

**Table 2:** Comparison of biofilm characteristics (1 *µ*m scalebar).

| Sample | Biofilm Area ($\mu\text{m}^2$) | Biofilm Coverage (%) |
| --- | --- | --- |
| **T1** | 4.12 | 24.4 |
| **T2** | 2.93 | 11.1 |
| **T3** | 4.65 | 19.5 |

## 4. Discussion

Injection of planktonic *B. subtilis* bacteria into the ECM of growing liquid cultures of *H. erinaceus* results in enveloping of some fraction of bacteria in the ECM and attachment to hyphae. The fungal origin of the ECM may be assumed due to experimental parameters including submerged liquid culture growth methods, use of EPS forming cultures and confirmation of formation of EPS materials in *H. erinaceus* liquid cultures prior to injection. The general tendency for wild-type mycelium to form EPS in liquid cultures should also be noted for cases where sealed growth chambers are used as in this study. However, the presence of connective structures between bacteria as well as connecting bacteria to hyphae suggest that co-production of EPS materials may have occured. Bacterial specific structures within a mixed biofilm cannot be ruled out.

Infection by bacteria either did not transpire at the timescale of the duration of our experiments or was not a specific aspect of the interaction. The orientation of the bacteria attached to the hyphae does not support a hypothesis of endofungal infection compared with previous SEM studies of known endofungal bacterial pathogens. Establishing mutualistic interactions or symbiotic relationships is, however, not possible within the constraints of this study. Lack of infection by bacteria as a single factor may not be indicative of mutualistic interactions. Another possibility,yet not directly evidenced here, is the production of bacterial nanostructures for nutrient transport [21] in combination with fungal biofilm formation. Further studies would need to confirm whether the attachment structures have any functional role beyond mechanical anchoring in order to provide clarity on these issues. Fourier transform infrared microspectroscopy (FTIR) would provide details to distinguish the molecular composition of the attachment structures. Imaging at different timescales of growth in combination with the aforementioned methods would also greatly enhance the conclusions of this study.

SEM imaging appears to be a viable method to investigate morphological features of co-cultures of bacteria and mycelia. Dehydration artifacts impact hyphal structures the most but can be limited by reducing the number of HDMS rinses. Modification of the ethanol series percentages may also be beneficial to the hydration of the hyphae. The attachment structures anchoring the bacteria to the hyphae were preserved along with the overall morphology of the bacteria. The overall structure of the fungal biofilm was not preserved at the macroscale (10-100 *µ*m). In the imaged samples hydration was preserved sufficiently to view bacterial interactions with mycelia in co-cultures where the mycelium is initially grown in liquid culture and forms a biofilm. For investigation of more specific features the co-cultures can be imaged at full hydration using environmental scanning electron microscopy [22]. However, the density of the mycelial biofilm may prohibit imaging of interior components including bacteria. This added opacity may reduce the visibility of the connective structures observed in this study. 3D scanning and slicing would improve these measurements taking into account the overall colony densities. However, this kind of measurement was not possible using the preparation methods discussed above. Epoxy-based encapsulation and slicing of the resulting samples could improve understanding of 3D details in future work. Brightfield microscopy techniques have intermittent issues in identifying co-culture features without the use of specifically targeted genetic and/or fluorescent probes [21].

The attachment structures may be relevant to our particular interest in the integration of multi-scalar extracellular electrical signaling in co-cultures. Biofilm components may facilitate electrical signaling modalities of the bacteria and fungi if they are present. Attachment of bacteria to fungal hyphae may have many downstream modulating effects including modification of bioelectrical states in the area local to the structures [23]. Both microelectrode array (MEA) and electrochemical impedance spectroscopy measurements can be used, among other methods, to investigate this modulating capacity over varying experimental timescales.

## 5. Conclusion

In this paper, we outlined an approach to identifying a connective morphological feature of biofilms in co-cultures of fungi and bacteria. Mycelium was grown in liquid cultures and planktonic bacteria was injected into the extracellular matrix of the mycelium. A single morphological structure, namely the matrix components connecting bacteria and hyphae, appeared to differentiate these cultures from trivial stochastic mixing of bacteria and mycelium in liquids.

## 6. Acknowledgement

The authors would like to thank David Patton for assistance with scanning electron microscopy. The research has been conducted under the framework of the FUNGATERIA (www.fungateria.eu) project, which has received funding from the European Union’s HORIZON-EIC-2021-PATHFINDER CHALLENGES programme under grant agreement No. 101071145. It is co-funded by the UK Research and Innovation grant No. 10048406.

## 7. Data availability

The datasets used and/or analysed during the current study available from the corresponding author on reasonable request.

---

**Figure 1:**
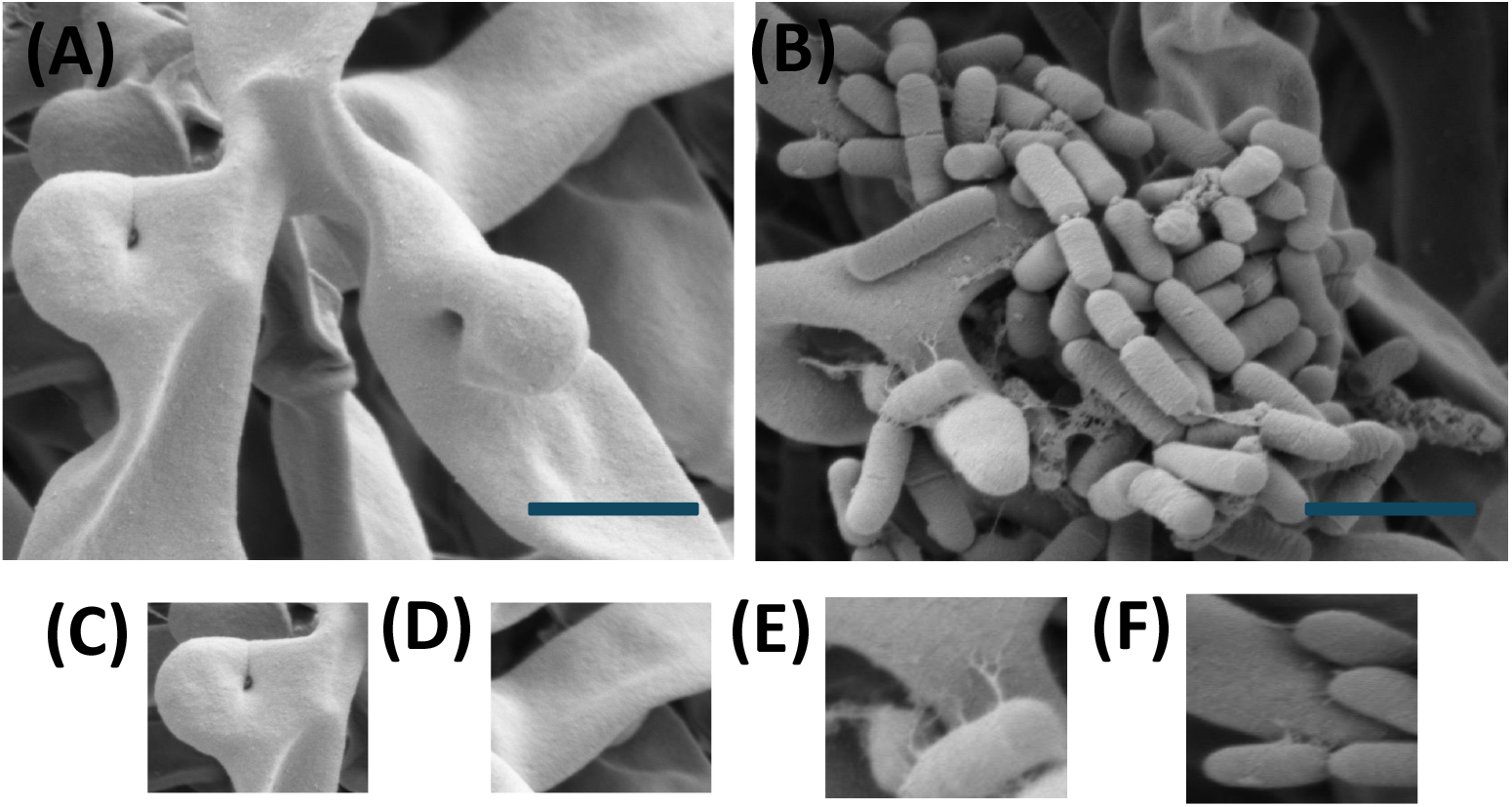
SEM micrographs of:(A) Mycelial monocultures (5 *µ*m scalebar, magnification of x9652, Hv of 2.00 kV, internal pressure of 2.36 × 10^−6^ Torr and HFW of 21.6 *µ*m);(B) Co-cultures (5 *µ*m scalebar, magnification of x8052, Hv of 2.00 kV, internal pressure of 4.93×10^−6^ Torr and HFW of 25.7 *µ*m);(C) Inset showing hyphal branching in mycelial monoculture samples;(D) Inset showing hyphal surface in mycelial monoculture samples;(E) Inset showing the attachment of the bacteria to hyphae via biofilm structures;(F) Inset showing the attachment of the bacteria to hyphae via biofilm structures.

**Figure 2:**
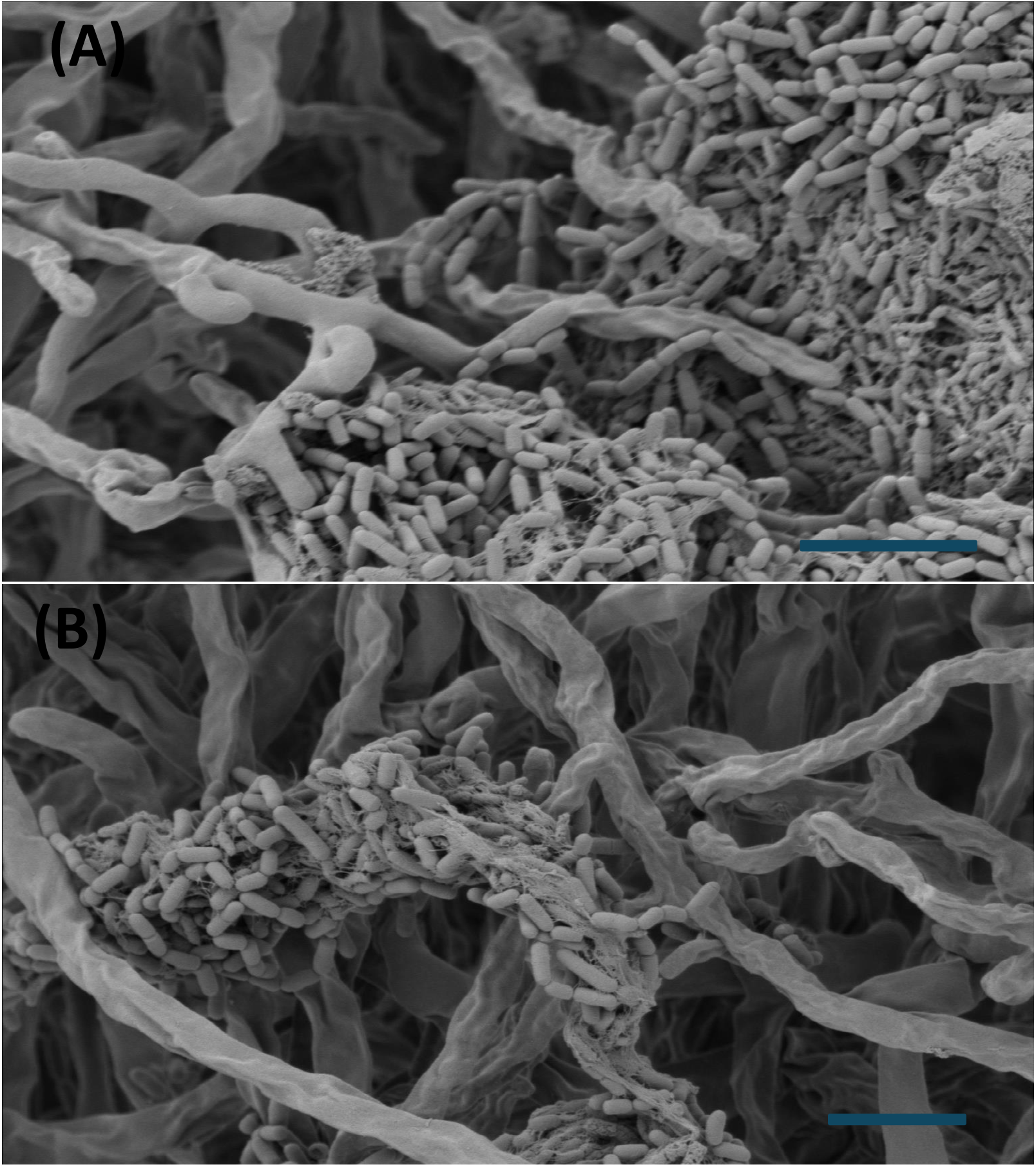
SEM micrographs of: (A) Co-cultures (5 *µ*m scalebar, magnification of x5572, Hv of 2.00 kV, internal pressure of 4.03×10^−6^ Torr and HFW of 37.2 *µ*m); (B) Co-cultures (5 *µ*m scalebar, magnification of x5453, Hv of 2.00 kV, internal pressure of 4.03×10^−6^ Torr and HFW of 38.0 *µ*m). Bacteria clustering on a single hyphae is a notable aspect of this micrograph.

**Figure 3:**
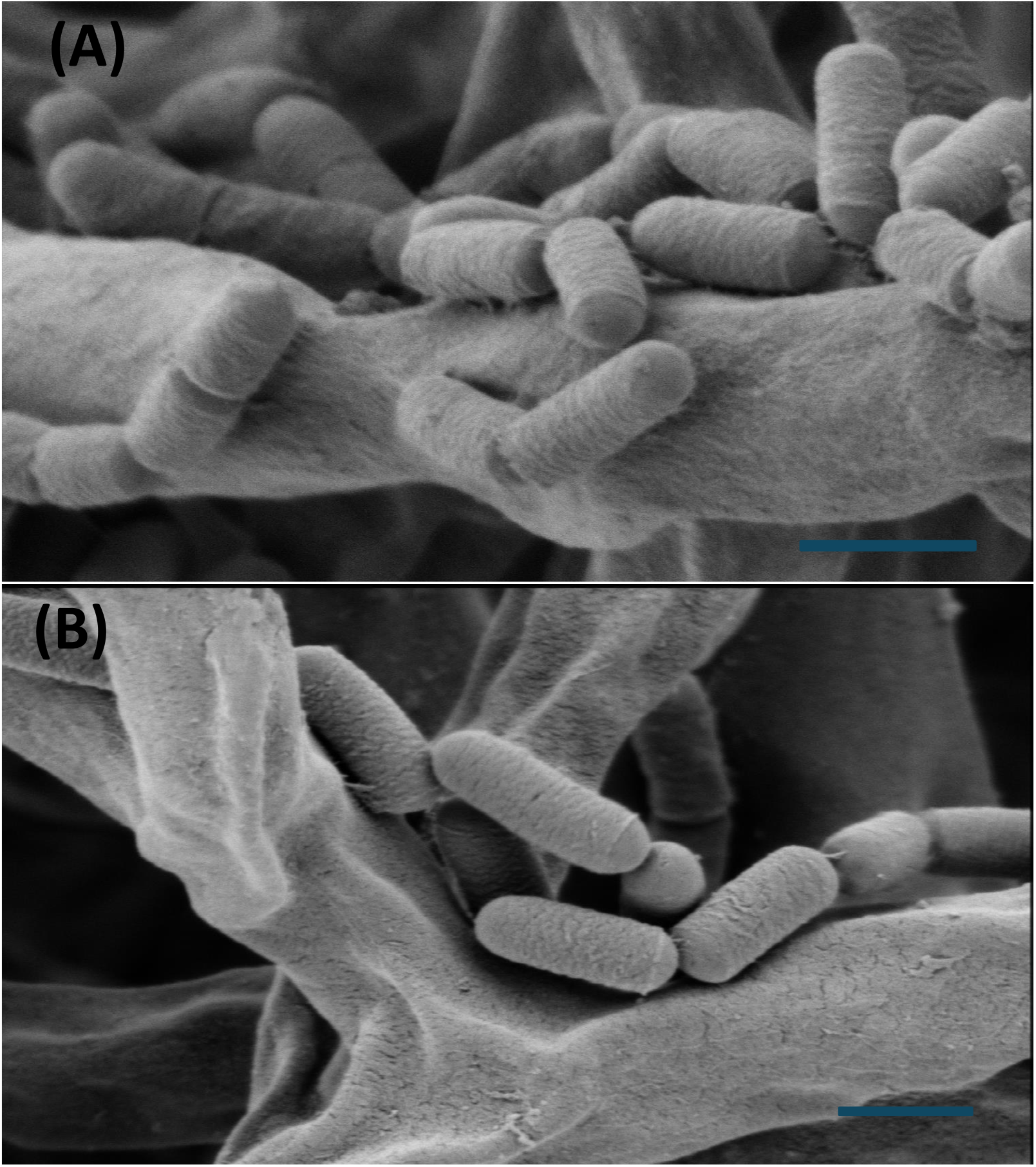
SEM micrographs of: (A) Co-cultures (1 *µ*m scalebar, magnification of x30000, Hv of 2.00 kV, internal pressure of 7.73×10^−6^ Torr and HFW of 7.65 *µ*m);(B) Co-cultures (1 *µ*m scalebar, magnification of x25481, Hv of 2.00 kV, internal pressure of 7.23×10^−6^ Torr and HFW of 8.13 *µ*m).

**Figure 4:**
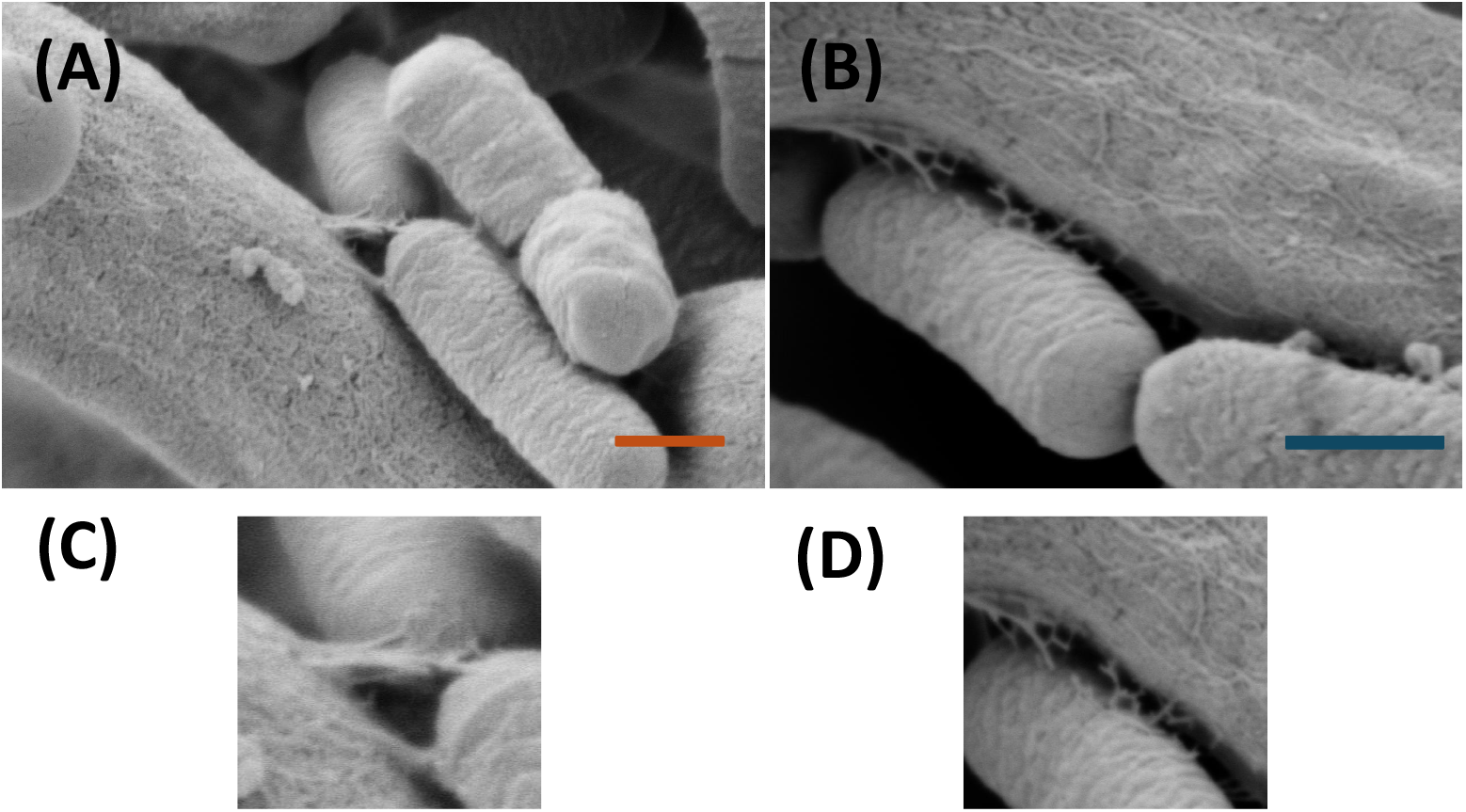
SEM micrographs of:(A) Co-cultures (500 *n*m scalebar, magnification of x51368, Hv of 2.30 kV, internal pressure of 3.44×10^−6^ Torr and HFW of 4.03 *µ*m);(B) Co-cultures (500 *n*m scalebar, magnification of x32428, Hv of 2.30 kV, internal pressure of 3.21×10^−6^ Torr and HFW of 6.39 *µ*m);(C) Zoomed inset of region of interest in (A) showing attachment structures; (D) Zoomed inset of region of interest in (B) showing attachment structures.

**Figure 5:**
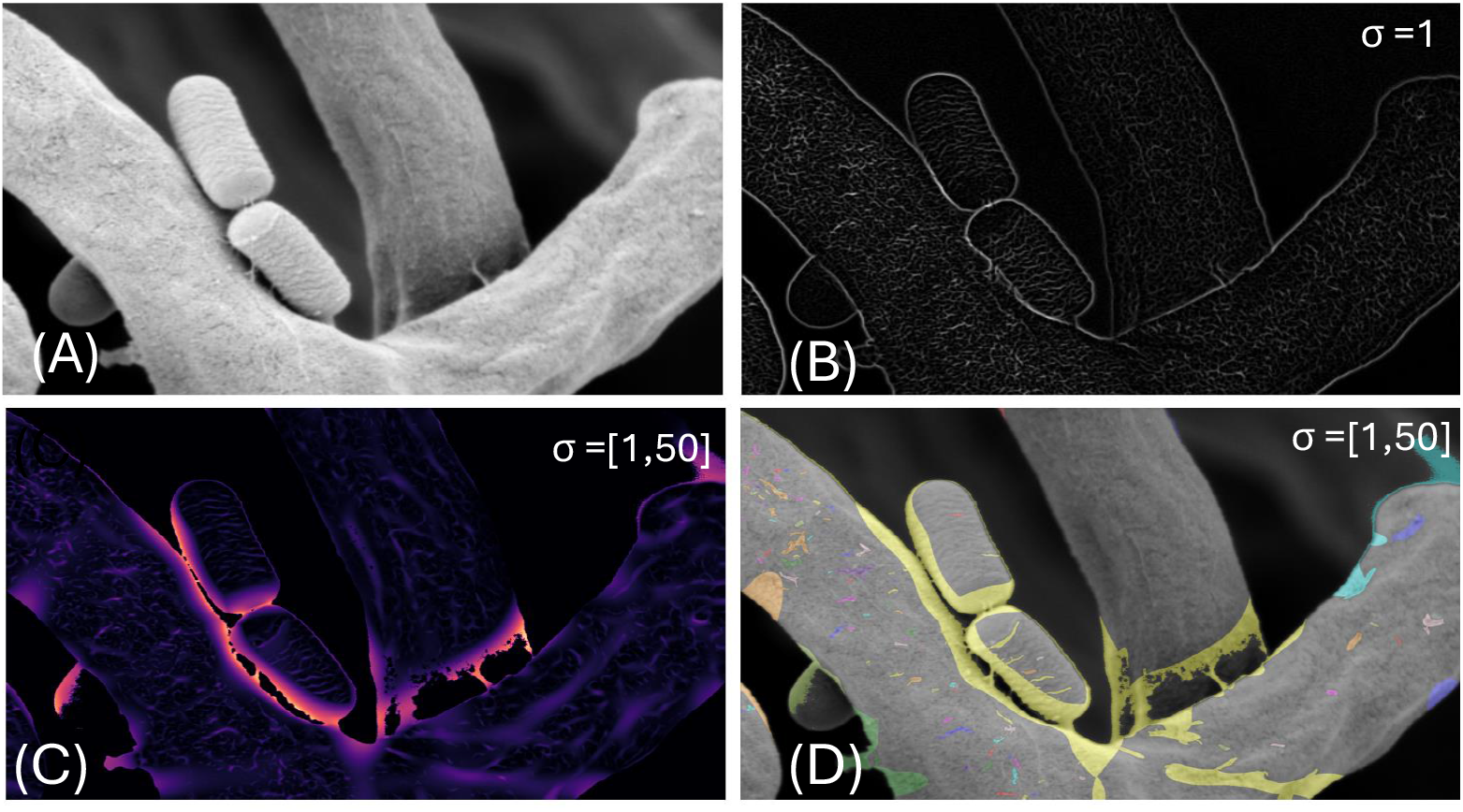
(A) Original micrograph;(B) Sato filter applied (*σ* = 1);(C) Identification of extracellular components distinct from the mycelia and bacteria using the Sato filter (*σ* = [1,50]);(D) Segmentation of biofilm components from fungal and bacterial structures based on the Sato filtering.

## References

[1] A. B. de Menezes, A. E. Richardson, P. H. Thrall, Linking fungal–bacterial co-occurrences to soil ecosystem function, Current Opinion in Microbiology 37 (2017) 135–141.

[2] F. Getzke, T. Thiergart, S. Hacquard, Contribution of bacterial-fungal balance to plant and animal health, Current opinion in microbiology 49 (2019) 66–72.

[3] P. Frey-Klett, P. Burlinson, A. Deveau, M. Barret, M. Tarkka, A. Sarniguet, Bacterial-fungal interactions: hyphens between agricultural, clinical, environmental, and food microbiologists, Microbiology and molecular biology reviews 75 (4) (2011) 583–609.

[4] W. Krüger, S. Vielreicher, M. Kapitan, I. D. Jacobsen, M. J. Niemiec, Fungal-bacterial interactions in health and disease, Pathogens 8 (2) (2019) 70.

[5] E. J. Espinosa-Ortiz, E. R. Rene, R. Gerlach, Potential use of fungal-bacterial co-cultures for the removal of organic pollutants, Critical reviews in biotechnology 42 (3) (2022) 361–383.

[6] H. Wilssens, L. De Wannemaeker, M. De Mey, Viable co-cultivation of filamentous fungi and engineered bacteria for fully biological and tuneable living materials, in: 7th Applied Synthetic Biology in Europe, 2024.

[7] A. Deveau, G. Bonito, J. Uehling, M. Paoletti, M. Becker, S. Bindschedler, S. Hacquard, V. Herve, J. Labbé, O. A. Lastovetsky, et al., Bacterial–fungal interactions: ecology, mechanisms and challenges, FEMS microbiology reviews 42 (3) (2018) 335–352.

[8] N. Moebius, Z. Üzüm, J. Dijksterhuis, G. Lackner, C. Hertweck, Active invasion of bacteria into living fungal cells, Elife 3 (2014) e03007.

[9] J. P. Shaffer, M. E. Carter, J. E. Spraker, M. Clark, B. A. Smith, K. L. Hockett, D. A. Baltrus, A. E. Arnold, Transcriptional profiles of a foliar fungal endophyte (pestalotiopsis, ascomycota) and its bacterial symbiont (luteibacter, gammaproteobacteria) reveal sulfur exchange and growth regulation during early phases of symbiotic interaction, Msystems 7 (2) (2022) e00091–22.

[10] G. H. Giger, C. Ernst, I. Richter, T. Gassler, C. M. Field, A. Sintsova, P. Kiefer, C. G. Gäbelein, O. Guillaume-Gentil, K. Scherlach, et al., Inducing novel endosymbioses by implanting bacteria in fungi, Nature (2024) 1–8.

[11] T. Hover, T. Maya, S. Ron, H. Sandovsky, Y. Shadkchan, N. Kijner, Y. Mitiagin, B. Fichtman, A. Harel, R. M. Shanks, et al., Mechanisms of bacterial (serratia marcescens) attachment to, migration along, and killing of fungal hyphae, Applied and Environmental Microbiology 82 (9) (2016) 2585–2594.

[12] L. C. Hsu, J. Fang, D. A. Borca-Tasciuc, R. W. Worobo, C. I. Moraru, Effect of micro-and nanoscale topography on the adhesion of bacterial cells to solid surfaces, Applied and environmental microbiology 79 (8) (2013) 2703–2712.

[13] R. Breitenbach, R. Gerrits, P. Dementyeva, N. Knabe, J. Schumacher, I. Feldmann, J. Radnik, M. Ryo, A. A. Gorbushina, The role of extracellular polymeric substances of fungal biofilms in mineral attachment and weathering, npj Materials degradation 6 (1) (2022) 42.

[14] S. Sieuwerts, F. A. De Bok, J. Hugenholtz, J. E. van Hylckama Vlieg, Unraveling microbial interactions in food fermentations: from classical to genomics approaches, Applied and environmental microbiology 74 (16) (2008) 4997–5007.

[15] E. Addis, G. Fleet, J. Cox, D. Kolak, T. Leung, The growth, properties and interactions of yeasts and bacteria associated with the maturation of camembert and blue-veined cheeses, International journal of food microbiology 69 (1-2) (2001) 25–36.

[16] M. Gou, Y. Qu, J. Zhou, F. Ma, L. Tan, Azo dye decolorization by a new fungal isolate, penicillium sp. qq and fungal-bacterial cocultures, Journal of Hazardous Materials 170 (1) (2009) 314–319.

[17] A. A. Brakhage, V. Schroeckh, Fungal secondary metabolites–strategies to activate silent gene clusters, Fungal Genetics and Biology 48 (1) (2011) 15–22.

[18] O. Tsivileva, A. Shaternikov, E. Ponomareva, Edible mushrooms could take advantage of the growth-promoting and biocontrol potential of, in: Proceedings of the Latvian Academy of Sciences. Section B. Natural, Exact, and Applied Sciences., Vol. 76, 2022, pp. 211–217.

[19] C. A. Schneider, W. S. Rasband, K. W. Eliceiri, Nih image to imagej: 25 years of image analysis, Nature methods 9 (7) (2012) 671–675.

[20] S. Van der Walt, J. L. Schönberger, J. Nunez-Iglesias, F. Boulogne, J. D. Warner, N. Yager, E. Gouillart, T. Yu, scikit-image: image processing in python, PeerJ 2 (2014) e453.

[21] E. Angulo-Cánovas, A. Bartual, R. López-Igual, I. Luque, N. P. Radzinski, I. Shilova, M. Anjur-Dietrich, G. García-Jurado, B. Úbeda, J. A. González-Reyes, et al., Direct interaction between marine cyanobacteria mediated by nanotubes, Science Advances 10 (21) (2024) eadj1539.

[22] G. Danilatos, Foundations of environmental scanning electron microscopy, in: Advances in electronics and electron physics, Vol. 71, Elsevier, 1988, pp. 109–250.

[23] T. Monk, N. Dennler, N. Ralph, S. Rastogi, S. Afshar, P. Urbizagastegui, R. Jarvis, A. van Schaik, A. Adamatzky, Electrical signaling beyond neurons, Neural Computation 36 (10) (2024) 1939–2029.

